# Representational shifts from feedforward to feedback rhythms index phenomenological integration in naturalistic vision

**DOI:** 10.1101/2024.09.17.613416

**Authors:** Lixiang Chen, Radoslaw Martin Cichy, Daniel Kaiser

## Abstract

How does the brain integrate complex and dynamic visual inputs into phenomenologically seamless percepts? Previous results demonstrate that when visual inputs are organized coherently across space and time, they are more strongly encoded in feedback-related alpha rhythms, and less strongly in feedforward-related gamma rhythms. Here, we tested whether this representational shift from feedforward to feedback rhythms is linked to the phenomenological experience of coherence. In an EEG study, we manipulated the degree of spatiotemporal coherence by presenting two segments from the same video across visual hemifields, either synchronously or asynchronously (with a delay between segments). We asked participants whether they perceived the stimulus as coherent or incoherent. When stimuli were presented at the perceptual threshold (i.e., when the same stimulus was judged as coherent 50% of times), perception co-varied with stimulus coding across alpha and gamma rhythms: When stimuli were perceived as coherent, they were represented in alpha activity; when stimuli were perceived as incoherent, they were represented in gamma activity. Whether the same visual input is perceived as coherent or incoherent thus depends on representational shifts between feedback-related alpha and feedforward-related gamma rhythms.

## Introduction

The visual inputs we receive in real life consist of vast arrays of features scattered across space and dynamically evolving through time. Yet, we phenomenologically experience the world in a spatiotemporally seamless manner. How does the brain integrate the complex and ever-changing inputs across space and time?

Information redundancy in natural inputs may play a critical role: Inputs are redundant across space, with predictable arrangements of both low-level features (Simoncelli and Olshausen, 2001) and high-level object content (Kaiser et al., 2019a). They are also redundant across time, with events unfolding in highly predictable sequences (Hogendoorn, 2022). These redundancies enable the brain to efficiently predict how visual features need to be integrated across space and time.

Such predictions are carried by cortical feedback flows expressed in dedicated rhythmic channels (van Kerkoerle et al., 2014; Michalareas et al., 2016): When natural inputs align across space and time, and thus can be integrated into a coherent percept, cortical feedback traverses the hierarchy in alpha rhythms from high-level visual cortex to early visual cortex, whereas processing is dominated by feedforward-related gamma activity when the inputs do not match up (Chen et al., 2023).

If rhythmic feedback was indeed critical for visual integration, the degree of feedback should covary with the phenomenological experience of a coherent visual world: When a visual input is perceived as coherent, it should be represented more strongly in feedback-related alpha rhythms, while inputs perceived as incoherent should be represented more strongly in feedforward-related gamma rhythms.

Here, we put this prediction to the test. In an EEG study, we presented natural video segments across the two visual hemifields, either synchronously or asynchronously (with one segment relatively delayed in time), and asked participants to report whether they perceived the stimulation as spatiotemporally coherent or not. Critically, this paradigm allowed us to test whether asynchronously presented videos at the perceptual threshold (i.e., stimuli perceived as coherent 50% of times) are coded differently across alpha and gamma rhythms, depending on the perceptual report.

## Results

We manipulated the degree of spatiotemporal coherence by presenting two segments from the same video synchronously or asynchronously (Fig. 1A) through two square apertures left and right of the central fixation (Fig. 1B). Twenty-six participants reported whether the whole video display appeared as coherent or incoherent to them. When temporal delays were low, the videos should appear as coherent (i.e., stemming from one seamless movie), but with greater delays they should appear as incoherent (i.e., with noticeable offset). To investigate differences in neural processing at the perceptual threshold, we initially quantified the delays that led to coherent and incoherent perception with equal probability separately for each of the five videos in a behavioral experiment (Fig. 1C).

**Figure 1.**
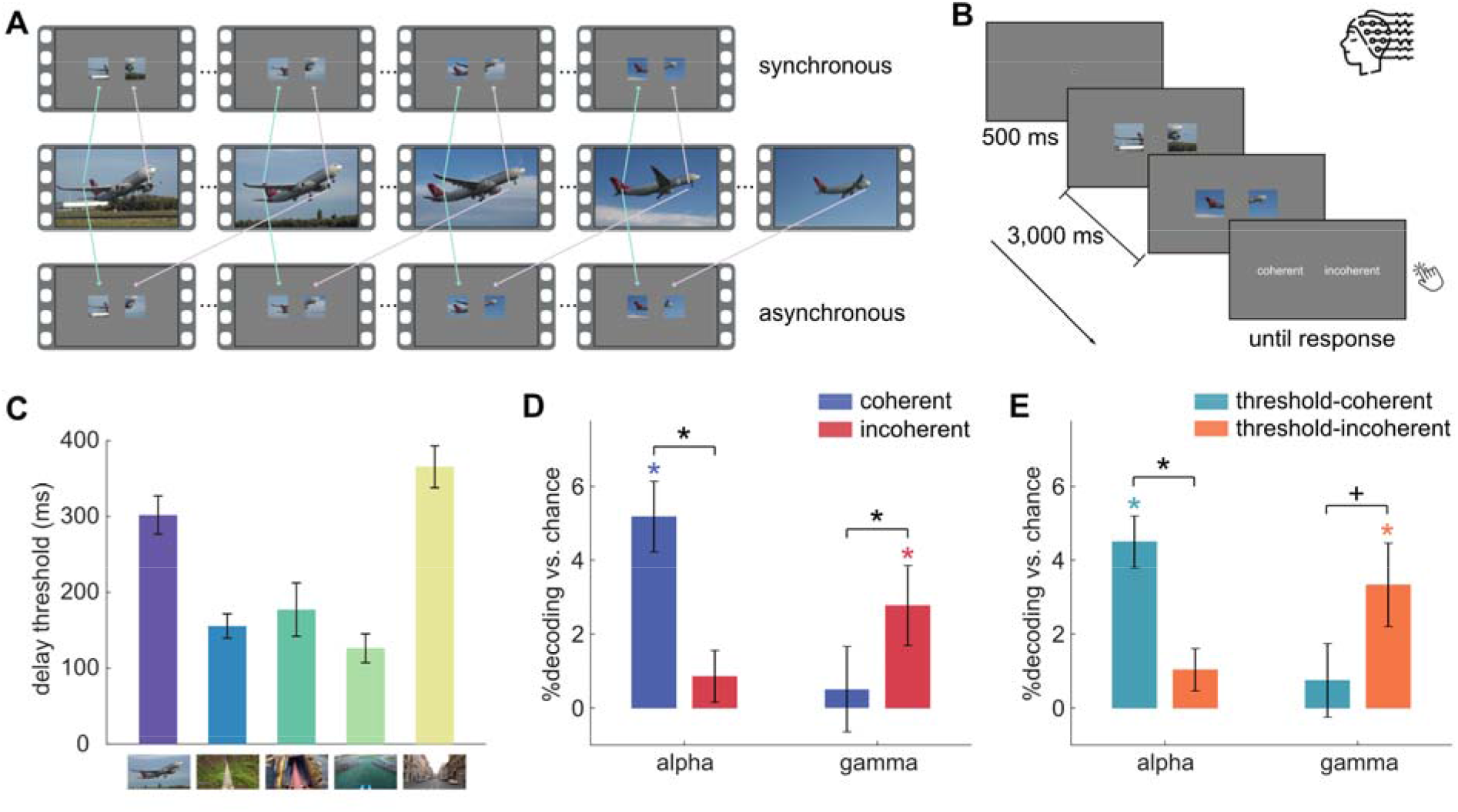
A rhythmic signature of phenomenological coherence. **(A)** Five natural videos were presented through apertures left and right of fixation, either synchronously or asynchronously. **(B)** Participants were instructed to fixate centrally and judge whether the stimulation was perceptually coherent or incoherent. **(C)** In an initial behavioral experiment, we used adaptive staircases to determine participants’ threshold delay for each of the five videos. In the subsequent EEG experiment, we presented the videos with no delay (coherent), the staircased delay (threshold), and twice the staircased delay (incoherent). We further separated the threshold trials into coherent and incoherent trials based on participants’ responses. We then decoded between the 5 videos within each condition using spectral power patterns in the alpha and gamma bands. **(D)** In the coherent and incoherent conditions, we found that coherent stimuli were decodable from alpha activity, suggesting prominent feedback propagation, whereas incoherent stimuli were decodable from gamma activity, suggesting dominant feedforward propagation. **(E)** The threshold condition replicated these results, showing that the representational balance across alpha and gamma rhythms tracks perceived coherence for identical visual inputs. Error bars represent standard errors. \**P* < 0.05, +*P* = 0.064.

In the subsequent EEG experiment, we presented the video stimuli in three conditions: no delay between the two video segments (coherent; 25% of trials), the delay at each participant’s subjective threshold (threshold; 50% of trials), and twice this subjective threshold (incoherent; 25% of trials). We subsequently split the threshold trials into threshold-coherent and threshold-incoherent trials, based on participants’ responses, allowing us to quantify neural representations for the same stimulus when participants perceived it as coherent or incoherent.

We hypothesized that the videos were coded more strongly in feedback-related alpha when perceived as coherent and more strongly in feedforward-related gamma when perceived as incoherent (Chen et al., 2023). To test this prediction, we decoded the five video stimuli in each condition using spectral power patterns across parietal-occipital channels, separately for the alpha (8–12 Hz) and gamma (31–70 Hz) frequency bands.

We found that stimuli in the coherent condition were decodable from alpha activity [*t*(25) = 5.409, *P* < 0.001], whereas stimuli in the incoherent condition were decodable from gamma activity [*t*(25) = 2.558, *P* = 0.017]. Coherent stimuli were decoded better than incoherent stimuli in the alpha frequency band [*t*(25) = 4.257, *P* < 0.001], and incoherent stimuli were decoded better than coherent stimuli in the gamma frequency band [*t*(25) = 2.203, *P* = 0.037; interaction: *F*(1, 25) = 30.282, *P* < 0.001] (Fig. 1D).

The threshold condition replicated this pattern of results: When the stimuli at threshold were perceived as coherent, they were decodable in the alpha band [*t*(25) = 6.415, *P* < 0.001], and when the stimuli were perceived as incoherent, they were decodable in the gamma band [*t*(25) = 2.948, *P* = 0.014]. Coherent stimuli were decoded better than incoherent stimuli in the alpha frequency band [*t*(25) = 5.385, *P* < 0.001], and incoherent stimuli were decoded better than coherent stimuli in the gamma frequency band, though only marginally [*t*(25) = 1.938, *P* = 0.064; interaction: *F*(1, 25) = 23.592, *P* < 0.001] (Fig. 1E).

No effects were found in the theta (4–7 Hz) and beta (13–30 Hz) frequency bands and in evoked broadband responses.

## Discussion

Our results show that the representational balance between alpha and gamma rhythms tracks the phenomenological experience of coherence: The same stimulus is coded in feedback-related alpha when it is perceived as coherent but in feedforward-related gamma when it is perceived as incoherent.

Integration-related alpha dynamics may carry spatiotemporally redundant – and thus predictable – stimulus information upstream (Bastos et al., 2012; van Kerkoerle et al., 2014; Michalareas et al., 2016), guiding the adaptive integration of this information into meaningful unified percepts.

We have previously demonstrated that spatiotemporally coherent stimulation, which readily allows for integration into a coherent percept, is linked to content coding in feedback-related alpha rhythms (Chen et al., 2023, 2024). Our new results show that these alpha-rhythmic codes indeed relate to the phenomenological experience of visual coherence.

Alpha rhythms have been associated with temporal integration before. The duration of the alpha cycle has been linked to the width of temporal integration windows (Samaha and Postle, 2015; VanRullen, 2016), and the phase and power of pre-stimulus alpha rhythms have been linked to integration versus segregation in subsequently presented stimuli (Leonardelli et al., 2015; Samaha and Postle, 2015). Our findings demonstrate that, beyond that, alpha rhythms also fulfill a function in representing the contents of upstream flows in cortex, suggesting an active involvement of alpha in binding visual stimuli across time (and space).

Our study probed the concurrent integration across space and time, and perceived incongruencies could originate from incongruent temporal patterns (e.g., motion trajectories) or spatial patterns (e.g., continuation of contours). Whether spatial or temporal properties drive integration to different extents needs to be explored further. Alternatively, integration across space or time may be governed by shared neural mechanisms (Walsh, 2003).

To conclude, our results suggest that representational shifts from bottom-up gamma to top-down alpha dynamics drive visual integration, highlighting the crucial role of cortical feedback in the construction of seamless perceptual experiences. More broadly, our results provide a rhythmic signature of the feedforward-to-feedback balance in visual cortex, which can be employed to track subjective changes in perception, attention, or cognition as a function of top-down or bottom-up dominance (Herz et al., 2020).

## Materials and Methods

### Participants

Twenty-six healthy adults (16 females; age = 22.2 ± 2.6 years) with normal or corrected-to-normal vision participated. A minimum sample size of 24 was determined with an effect size of 0.25 as derived from our previous study (Chen et al., 2023), a significance level of 0.05, and a power of 0.8. All participants signed written informed consent and received either course credits or cash reimbursement. The study was approved by the ethical committee of the Department of Education and Psychology at Freie Universität Berlin and was conducted in accordance with the Declaration of Helsinki.

### Stimuli and Paradigm

The stimulus set consisted of five short video clips (airplane takeoff, cyclist, roller coaster, ski jumper, and driving car). The videos were presented through two square apertures (6° visual angle) left and right of the central fixation (2.78° offset). The central fixation had a diameter of 0.44° visual angle. Videos were played either synchronously (in each frame, the two images shown through the apertures were from the same frame of the original video) or asynchronously (in each frame, the two images were from different frames of the original video; see Fig. 1A). We presented stimuli (at 60-Hz refresh rate) and recorded participants’ responses using MATLAB and the Psychophysics Toolbox (Brainard, 1997). We first presented a central fixation for 0.5 seconds, followed by the videos for 3 seconds. Participants were instructed to maintain central fixation and, after the video ended, judge whether the videos were perceptually coherent or incoherent. An example trial is shown in Fig. 1B.

### Behavioral Experiment

We first conducted a behavioral experiment to estimate subjective integration thresholds for scenes using the QUEST adaptive staircasing procedure (Watson and Pelli, 1983). We ran separate QUEST staircases for each video, initializing the delay between two video segments randomly between 100 and 400 ms in the first trial of each scene and adaptively adjusting the delay afterwards. Each staircase terminated after 80 trials. The staircases converged within this trial count. For each participant, we averaged the delay values in the last 5 trials for each video to obtain the threshold delays.

### EEG Experiment

In the EEG experiment, we presented stimuli in three conditions. In the coherent condition, stimuli were presented synchronously. In the threshold condition, we set the delay between video segments for each scene to the subjective threshold estimated in the behavioral experiment with the same participant. In the incoherent condition, we set the delay for each scene to twice the subjective threshold. Each coherent/incoherent stimulus was presented 30 times, and each threshold stimulus was shown 60 times, yielding 600 trials, which were presented in random order. For the conditions with a delay, the left segment temporally led in half of the trials, and the right segment led in the other half. After the experiment, we separated threshold trials based on each participant’s responses: if a trial was judged as coherent, we assigned it to the threshold-coherent condition; otherwise, we assigned the trial to the threshold-incoherent condition. We recorded EEG and eye-tracking data while participants conducted the experiment. EEG data were acquired using a 10-10 EASYCAP 64-electrode system with a BrainVision actiCHamp amplifier at 1000 Hz. The data were online filtered at 0.03–100 Hz and referenced to FCz. Eye-tracking data were acquired using the Psychophysics and Eyelink Toolbox extensions (Cornelissen et al., 2002), with an Eyelink 1000 Tower Mount (SR Research Ltd., Canada). We recorded movements of the right eye and conducted a standard 9-point calibration at the beginning of the experiment.

### Eye-Movement Analysis

We used Fieldtrip (Oostenveld et al., 2011) to epoch the eye-tracking data from -0.5 to 3.5 s and downsampled the data to 200 Hz. To check participants’ fixation stability, we estimated the mean and standard deviation (SD) of horizontal eye movements during stimulus presentation (0–3 seconds). There were no significant between-condition differences in horizontal eye movements, neither in the mean, *F*(3,75) = 1.295, *P* = 0.295, nor the standard deviation, *F*(3,75) = 0.334, *P* = 0.801.

### EEG Preprocessing

We preprocessed EEG data using Fieldtrip. We first epoched the data from -0.5 to 3.5 s relative to the stimulus onset. We then band-stop filtered the data to remove 50-Hz line noise, referenced the data to the average of all channels, and downsampled the data to 200 Hz. Next, we visually inspected the data and removed noisy trials and channels. The removed channels were subsequently interpolated using their neighboring channels. We performed independent component analysis to further remove blinks and eye movement artifacts. Finally, the data were baseline-corrected by subtracting the mean of pre-stimulus signals.

### EEG Spectral Analysis

We performed spectral analysis on the preprocessed EEG data using Fieldtrip, replicating the analysis pipeline used previously (Chen et al., 2023, 2024). For each trial, we conducted the fast Fourier transform (FFT) and estimated the power of each frequency from 4 to 70 Hz in each channel. We used a signal tapper with a Hanning window for the low-frequency bands: theta (4–7 Hz, in steps of 1 Hz), alpha (8–12 Hz, in steps of 1 Hz), and beta (13–30 Hz, in steps of 2 Hz). For the gamma band (31–70 Hz, in steps of 2 Hz), we used the discrete prolate spheroidal sequences (DPSS) multitaper method with ±8 Hz smoothing.

### EEG Decoding Analysis

We performed multivariate decoding analysis to probe rhythmic representations of stimuli using CoSMoMVPA (Oosterhof et al., 2016) and LIBSVM (Chang and Lin, 2011). For this analysis, we chose 17 parietal and occipital (PO) channels (Pz, P1, P2, P3, P4, P5, P6, P7, P8, POz, PO3, PO4, PO7, PO8, Oz, O1, O2) over visual cortex (Kaiser et al., 2019b; Chen et al., 2024) and extracted the spectral power patterns across these channels to differentiate between five stimuli within each of the four conditions (coherent, threshold-coherent, threshold-incoherent, and incoherent), separately for each frequency band (theta, alpha, beta, and gamma). The analysis was conducted using the linear support machine (SVM) and leave-one-trial-out cross-validation. The number of trials was always balanced across scenes. Additionally, to reduce the dimensionality of the data, we performed PCA on the training data and then projected the PCA solutions (99% variance explained of the training set) onto the testing data (Chen et al., 2022, 2023, 2024). We compared the decoding accuracy against the chance level (20%) separately for each frequency band and each condition to detect frequency-specific representations of stimuli. To investigate whether the representations are modulated by experimental conditions, we performed 2-condition × 4-frequency two-way ANOVAs separately for the coherent/incoherent and the threshold-coherent/threshold-incoherent conditions, and t-tests against chance level for each condition. Multiple comparisons were corrected using false discovery rate (FDR) correction (*P* < 0.05).

## Acknowledgments

L.C. is supported by a PhD stipend from the China Scholarship Council (CSC). R.M.C is supported by the Deutsche Forschungsgemeinschaft (DFG; CI241/3-1, and INST 272/297-1) and by a European Research Council (ERC) starting grant (ERC-2018-STG 803370). D.K. is supported by the DFG (SFB/TRR135, project number 222641018; KA4683/5-1, project number 518483074; KA4683/6-1, project number 536053998), “The Adaptive Mind” funded by the Excellence Program of the Hessian Ministry of Higher Education, Science, Research and Art, and an ERC Starting Grant (ERC-2022-STG 101076057). Views and opinions expressed are those of the authors only and do not necessarily reflect those of the European Union or the European Research Council. Neither the European Union nor the granting authority can be held responsible for them. The authors thank the HPC Service of ZEDAT, Freie Universität Berlin, for computing time.

## Author Contributions

L.C. and D.K. designed research; L.C. performed research; L.C. analyzed data; and L.C., D.K., and R.M.C. wrote the paper.

## References

Bastos AM, Usrey WM, Adams RA, Mangun GR, Fries P, Friston KJ (2012) Canonical microcircuits for predictive coding. Neuron 76:695–711.

Brainard DH (1997) The Psychophysics Toolbox. Spat Vis 10:433–436.

Chang C-C, Lin C-J (2011) LIBSVM: a library for support vector machines. ACM Trans Intell Syst Technol 2:1–27.

Chen L, Cichy RM, Kaiser D (2022) Semantic scene-object consistency modulates N300/400 EEG components, but does not automatically facilitate object representations. Cereb Cortex 32:3553–3567.

Chen L, Cichy RM, Kaiser D (2023) Alpha-frequency feedback to early visual cortex orchestrates coherent naturalistic vision. Sci Adv 9:eadi2321.

Chen L, Cichy RM, Kaiser D (2024) Coherent categorical information triggers integration-related alpha dynamics. J Neurophysiol 131:619–625.

Cornelissen FW, Peters EM, Palmer J (2002) The Eyelink Toolbox: Eye tracking with MATLAB and the Psychophysics Toolbox. Behav Res Methods Instrum Comput 34:613–617.

Herz N, Baror S, Bar M (2020) Overarching States of Mind. Trends Cogn Sci 24:184–199.

Hogendoorn H (2022) Perception in real-time: predicting the present, reconstructing the past. Trends Cogn Sci 26:128–141.

Kaiser D, Quek GL, Cichy RM, Peelen MV (2019a) Object vision in a structured world. Trends Cogn Sci 23:672–685.

Kaiser D, Turini J, Cichy RM (2019b) A neural mechanism for contextualizing fragmented inputs during naturalistic vision. Elife 8:e48182.

Leonardelli E, Braun C, Weisz N, Lithari C, Occelli V, Zampini M (2015) Prestimulus oscillatory alpha power and connectivity patterns predispose perceptual integration of an audio and a tactile stimulus. Hum Brain Mapp 36:3486–3498.

Michalareas G, Vezoli J, van Pelt S, Schoffelen J-M, Kennedy H, Fries P (2016) Alpha-Beta and Gamma Rhythms Subserve Feedback and Feedforward Influences among Human Visual Cortical Areas. Neuron 89:384–397.

Oostenveld R, Fries P, Maris E, Schoffelen J-M (2011) FieldTrip: open source software for advanced analysis of MEG, EEG, and invasive electrophysiological data. Comput Intell Neurosci 2011:156869.

Oosterhof NN, Connolly AC, Haxby JV (2016) CoSMoMVPA: multi-modal multivariate pattern analysis of neuroimaging data in Matlab/GNU Octave. Front Neuroinformatics 10:27.

Samaha J, Postle BR (2015) The Speed of Alpha-Band Oscillations Predicts the Temporal Resolution of Visual Perception. Curr Biol 25:2985–2990.

Simoncelli EP, Olshausen BA (2001) Natural image statistics and neural representation. Annu Rev Neurosci 24:1193–1216.

van Kerkoerle T, Self MW, Dagnino B, Gariel-Mathis M-A, Poort J, van der Togt C, Roelfsema PR (2014) Alpha and gamma oscillations characterize feedback and feedforward processing in monkey visual cortex. Proc Natl Acad Sci 111:14332–14341.

VanRullen R (2016) Perceptual Cycles. Trends Cogn Sci 20:723–735.

Walsh V (2003) A theory of magnitude: common cortical metrics of time, space and quantity. Trends Cogn Sci 7:483–488.

Watson AB, Pelli DG (1983) Quest: A Bayesian adaptive psychometric method. Percept Psychophys 33:113–120.

